# Characterizing the role of autaptic feedback in enhancing precision of neuronal firing times

**DOI:** 10.1101/2023.10.06.561207

**Authors:** Zahra Vahdat, Oliver Gambrell, Abhyudai Singh

## Abstract

In a chemical synapse, information flow occurs via the release of neurotransmitters from a presynaptic neuron that triggers an Action potential (AP) in the postsynaptic neuron. At its core, this occurs via the postsynaptic membrane potential integrating neurotransmitter-induced synaptic currents, and AP generation occurs when potential reaches a critical threshold. This manuscript investigates feedback implementation via an autapse, where the axon from the postsynaptic neuron forms an inhibitory synapse onto itself. Using a stochastic model of neuronal synaptic transmission, we formulate AP generation as a first-passage time problem and derive expressions for both the mean and noise of AP-firing times. Our analytical results supported by stochastic simulations identify parameter regimes where autaptic feedback transmission enhances the precision of AP firing times consistent with experimental data. These noise attenuating regimes are intuitively based on two orthogonal mechanisms - either expanding the time window to integrate noisy upstream signals; or by linearizing the mean voltage increase over time. Interestingly, we find regimes for noise amplification that specifically occur when the inhibitory synapse has a low probability of release for synaptic vesicles. In summary, this work explores feedback modulation of the stochastic dynamics of autaptic neurotransmission and reveals its function of creating more regular AP firing patterns.

## I. Introduction

Action potential (AP) triggered release of neurotransmitters from a presynaptic neuron to generate an AP in a postsynaptic neuron is a hallmark of interneuronal communication. There is a rich body of literature modeling the neurotransmitter release over time, including capturing noise mechanisms inherent in this process [1]–[8].

The simplest possible formulation of postsynaptic AP generation is the integrate-and-fire model [9], [10], where postsynaptic membrane potential increases in response to the neurotransmitter activity (assuming an excitatory synapse), and AP-firing occurs when the potential crosses a critical threshold. This model is often “leaky” which refers to the fact that voltage can decay back to resting potential when there is a lapse in upstream neurotransmitter activity.

Given that information is often encoded in AP frequency, how neurons ensure a precise firing pattern is an important and fundamental problem. Towards that end, we explore the role of a specific form of negative feedback known as an autapse (Fig. 1), where the postsynaptic neuron forms an inhibitory synapse on itself. Such autaptic transmission has been seen in neurons both in-culture and in-vivo [11]–[16]. A combination of computational and experimental works have implicated autapses in diverse functional roles from modulating the temporal fidelity of AP firing patterns [17]– [20], inducing oscillations [21]–[23], and synchronization of neuronal networks [24]–[26]. Our approach to modeling this system leverages the Stochastic Hybrid Systems (SHS) framework that couples continuous dynamical changes in variables (such as the membrane potential) with discrete events occurring at random times (such as AP firings). Building up on our previous work that mechanistically captures different noise sources in synaptic transmission as an SHS [27]–[31], we expand the model to now have two chemical synapses - an excitatory synapse from the presynaptic neuron and an inhibitory autapse (Fig. 1).

**Fig. 1.**
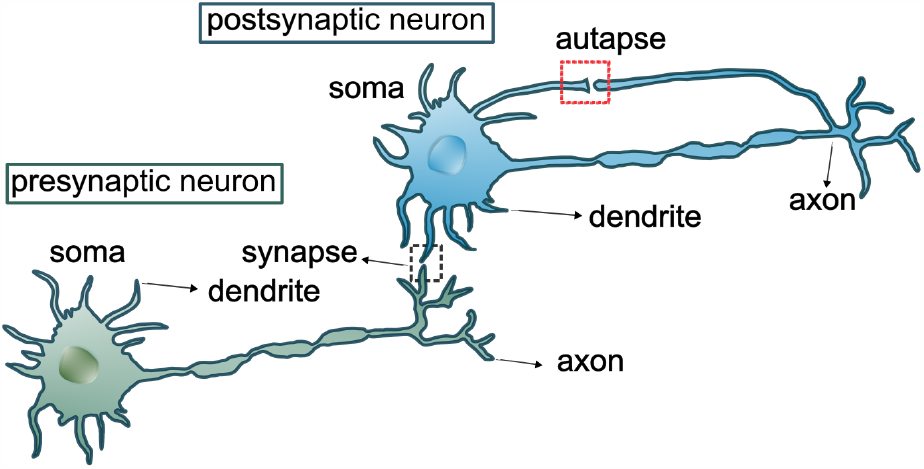
Schematic of a presynaptic neuron forming an excitatory synapse with a postsynaptic neuron. Autaptic feedback is implemented by having an axon of the postsynaptic neuron forming an inhibitory synapse on itself.

Formulating AP-triggering as a first-passage-time problem, we develop analytical formulas for the mean frequency of postsynaptic AP generation and the corresponding noise in firing times. The analytical approach identifies parameter regimes where an autaptic feedback – where a neuron forms an inhibitory synapse on itself – enhances AP firing precision and other regimes where noise is amplified making AP firings more irregular. Our analysis shows an interesting trade-off, where the negative feedback expands the time window for integrating upstream neurotransmitter release that attenuates noise. However, this comes at the cost of feedback itself introducing additional noise, as vesicle release in the inhibitory autapse is also a stochastic process. These results are confirmed by numerical simulations of the SHS.

## II. Formulating neurotransmission as a Stochastic Hybrid System

We start by reviewing the SHS formalism for capturing the timing of AP-generation in the postsynaptic neuron [27], [29], [30]. Later in this section, we extend the model to implement negative feedback via an autapse. For convenience, all model parameters are summarized in Table. I. Three stochastic processes are considered in the model-***n***(*t*) and ***n***_*d*_(*t*) are the number of vesicle-filled docking sites in the presynaptic and autaptic axon terminals at time *t*, respectively, and ***v***(*t*) is the postsynaptic membrane potential at time *t*. We denote random variables and stochastic processes by bold symbols.

**TABLE I.**
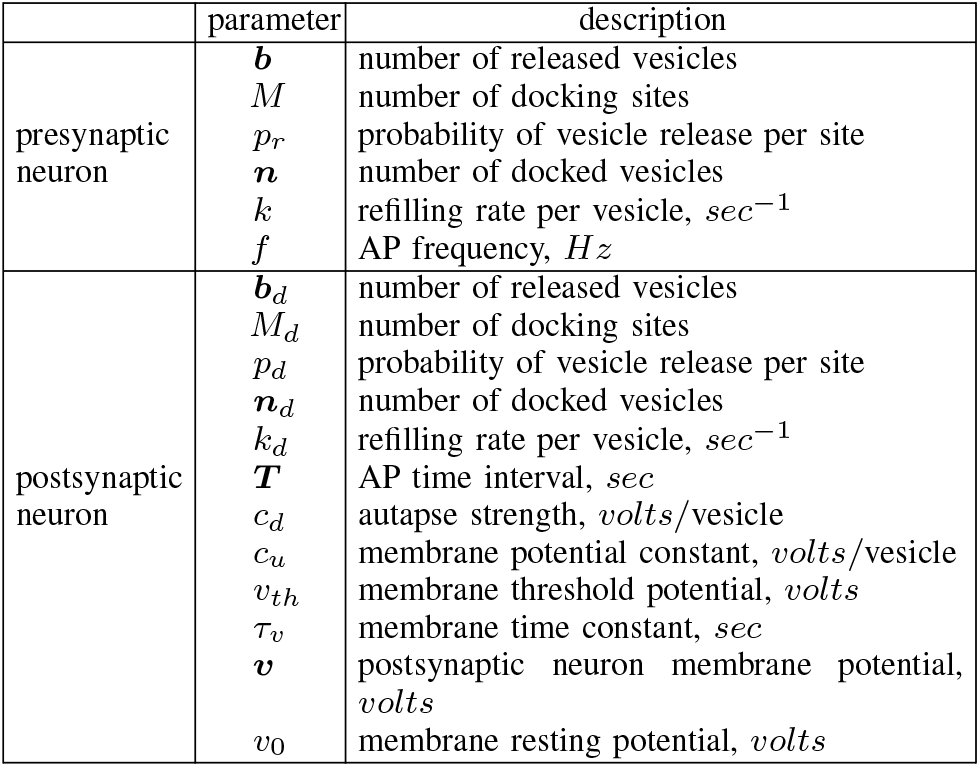
Summary of model parameters.

### A. Modeling presynaptic processes

Let APs be generated in the presynaptic neuron with frequency *f* as per a Poisson process, where the time between two successive APs is an exponentially-distributed random variable with mean 1*/f*. While our previous work generalized these results to consider any arbitrary inter-AP firing time distribution [27], [30], for simplicity we only consider the Poisson arrival case here. These APs travel to the active zone of the presynaptic axon terminal, which is assumed to have *M* ∈ { 1, 2, … } docking sites, and each site is empty or occupied by a synaptic vesicle that is filled with neurotransmitters.

Let ***n***(*t*) ∈ { 0, 1, …, *M*} denote the number of docking sites that are occupied with vesicles at time *t*. Whenever an AP arrives, each vesicle is released with probability *p*_*r*_ resulting in the reset

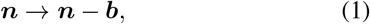

where ***b*** is drawn from a Binomial distribution

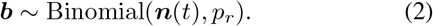

In between two consecutive APs, each empty docking site is replenished with a rate *k*. Given that there are *M* − ***n***(*t*) empty sites, the net replenishment rate is *k*(*M* − ***n***(*t*)). This stochastic replenishment process can be probabilistically defined as

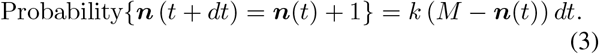

Combining (3) with (1), the two resets driving the stochastic dynamics of ***n***(*t*) can be represented together as

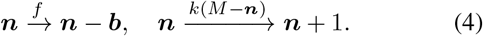

Using the standard tools of moment dynamics [32], we had previously shown that the first and second-order statistical moments of the scalar integer-valued random process ***n***(*t*) evolve as

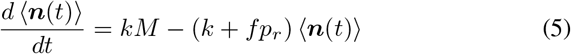

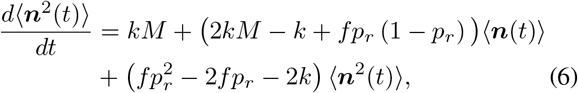

where the angular brackets ⟨ ⟩ denote the expected-value operation [27], [29]. Solving these equations at steady state yield

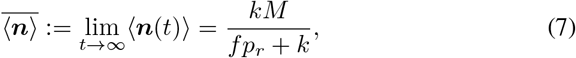

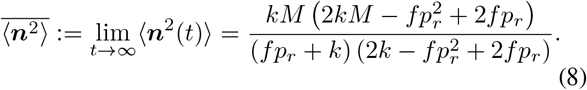

Our prior work has investigated the steady-state noise levels in both ***n***, and the number of released vesicles ***b***, as a function of model parameters *f, M, k, p*_*r*_ [27], [30], and also shown the utility of these results in the inference of parameters from experimental data [28].

### B. Modeling postsynaptic processes

Assuming rapid turnover of the released neurotransmitter, AP-triggered evoked release of vesicles results in an instantaneous jump in the postsynaptic membrane potential ***v***(*t*)

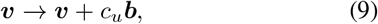

where *c*_*u*_ is a positive constant that is determined by a variety of factors, such as the number of neurotransmitters per vesicle, neurotransmitter binding kinetics to postsynaptic receptors, and the integration of induced synaptic currents. In between two presynaptic APs, the membrane potential decays exponentially as per first-order kinetics

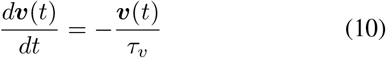

where *τ*_*v*_ is the membrane time constant and results in a “leaky” integrator. In the limit *τ*_*v*_ → ∞ this reduces to a pure integrator where ***v***(*t*) monotonically increases with time.

From a resting potential *v*_0_, ***v***(*t*) increases increases according to the jump-decay process (9)-(10). When ***v***(*t*) crosses a critical threshold *v*_*th*_, a postsynaptic AP is generated, and the potential resets back to *v*_0_.

Given that the membrane potential is a stochastic process, its mean levels evolve as

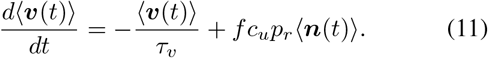

Moreover, the time evolution of second-order moments ⟨***v***^2^(*t*)⟩ and ⟨***n***(*t*)***v***(*t*)⟩ are given by

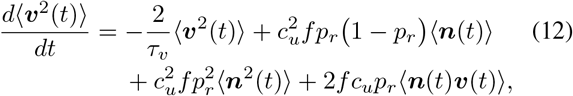

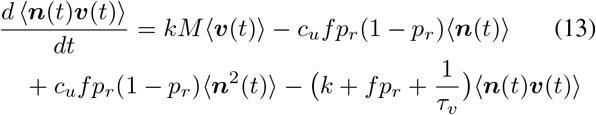

[33]. It is important to point out that the overall moment dynamics as given by (5)-(6), and (11)-(13) is a linear system that can be solved exactly for any arbitrary initial condition. We will leverage these solutions to study stochasticity in the timing of postsynaptic APs. However, before getting into this analysis, we discuss how autaptic feedback is implemented in the model.

### C. Implementing autaptic feedback

Autaptic feedback is implemented by having the axon from the postsynaptic neuron form an inhibitory synapse with itself (Fig. 1). We assume that the axon terminal of the postsynaptic neuron has *M*_*d*_ docking sites, ***n***_*d*_(*t*) sites are occupied with vesicles at time *t*, and each empty site fills with rate *k*_*d*_. The subscript *d* here denotes the processes and parameters in the postsynaptic neuron. Whenever a postsynaptic AP is generated, ***b***_*d*_ vesicles are released as per a Binomial distribution

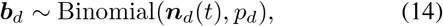

with *p*_*d*_ being the release probability per docked vesicle.

Analogous to (4), the stochastic dynamics of ***n***_*d*_(*t*) can be represented as

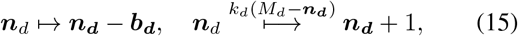

where the first reset occurs whenever there is a postsynaptic AP. Feedback results from the postsynaptic AP-triggered released neurotransmitter resetting its own neuron’s membrane potential as

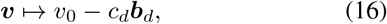

for some positive constant *c*_*d*_ that can be interpreted as the *autapse strength*. Thus, any postsynaptic AP firing will hyperpolarize the membrane potential below the resting potential *v*_0_ delaying the next AP-firing event.

It is important to point out that while the presynaptic-postsynaptic synapse is excitatory, the autapse is inhibitory, implying that both synapses use different types of neurotransmitters. The overall system with autaptic feedback can be represented as a SHS (Fig. 2A). The continuous dynamics are given by (10) and are depicted inside the oval. Different types of stochastic resets are indicated by arrows around the oval. The stochastic resets include

- Generation of APs in the presynaptic neuron, release of docked vesicles from the presynaptic axon terminal, postsynaptic membrane potential depolarization (jump to a larger value).
- Replenishment of empty sites in the presynaptic axon terminal.
- Release of docked vesicles from the postsynaptic axon terminal, postsynaptic membrane potential hyperpolarization (reset to resting potential and autaptic feedback effect) and AP triggering in the postsynaptic neuron.
- Replenishment of empty sites in the postsynaptic axon terminal.

**Fig. 2.**
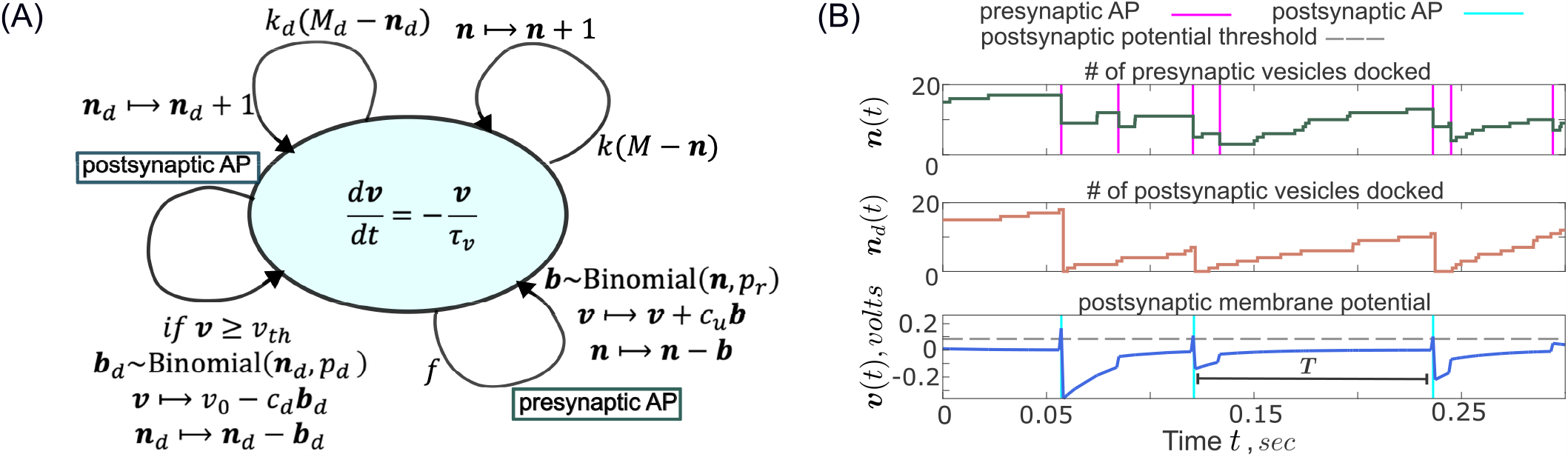
**A**. Model formulation of autaptic neurotransmission as a Stochastic Hybrid System (SHS) that consists of three random processes - ***n***(*t*) and ***n***_*d*_(*t*) are the number of vesicle-filled docking sites in the presynaptic and autaptic axon terminals, respectively, and ***v***(*t*) is the postsynaptic membrane potential. Presynaptic APs arrive with frequency *f* as per a Poisson process and postsynaptic APs are triggered when ***v***(*t*) reaches a threshold *v*_*th*_. See Section II for more SHS model details. **B**. Sample trajectories of the number of vesicles docked in the presynaptic axon terminal (top) and autaptic axon terminal (middle). APs trigger a release of vesicles in the respective neurons, and vesicles are replenished at a constant rate. The bottom plot is the postsynaptic membrane potential that results in a postsynaptic AP upon reaching the threshold potential. For these trajectories, vesicles docking rates are *k* = *k*_*d*_ = 10 *sec*^*−*1^ per vesicle, the membrane potential constant is *c*_*u*_ = .02 *volts/vesicle*, and the autapse strength is *c*_*d*_ = 0.02 *volts/vesicle*. Other parameters are taken as *f* = 30 *Hz, M* = *M*_*d*_ = 20, *p*_*r*_ = *p*_*d*_ = 0.4, *v*_*th*_ = 0.082 *volts, v*_0_ = 0.

A sample realization of the stochastic processes ***n***(*t*), ***n***_*d*_(*t*) and ***v***(*t*) is shown in Fig. 2B with AP generation events.

## III. Analysis of postsynaptic action potential time

We use the First Passage Time (FPT) concept to capture the time ***T*** between two successive postsynaptic APs. Mathematically, it can be defined as

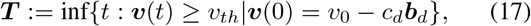

i.e., the first time the membrane potential reaches the threshold *v*_*th*_ starting from an initial voltage that is determined by (16). Here *c*_*d*_ = 0 corresponds to open-loop neurotransmission where voltage resets back to resting potential *v*_0_ after AP firing. Without loss of any generality, we assume *v*_0_ = 0.

Our goal is to investigate the mean postsynaptic AP firing times ⟨***T*** ⟩, and its noise level

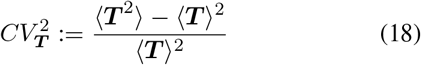

quantified through the dimensionless coefficient of variation squared 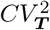. To obtain analytical insights we make approximations that will be subsequently relaxed by performing stochastic simulations of the full model. For example, considering *k*_*d*_→ ∞, i.e., an infinite vesicle refilling rate ensures all *M*_*d*_ docking sites in the postsynaptic terminal are filled with vesicles and available for release. This makes ***b***_*d*_∼ Binomial(*M*_*d*_, *p*_*d*_), and the voltage reset (16) independent and identically distributed random variables, and facilitates analytical derivation of closed-form formulas.

In the small-noise limit, the mean firing times ⟨ ***T*** ⟩ at steady-state can be simply obtained by solving

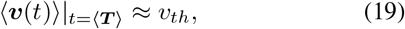

where ⟨***v***(*t*)⟩ is the solution to the first-order system (11) with initial condition ⟨***v***(0)⟩ = −*c*_*d*_*M*_*d*_*p*_*d*_ and ⟨***n***(*t*)⟩ replaced by 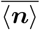 in (7). The mean voltage buildup is given by (32) in the Appendix. This approach yields

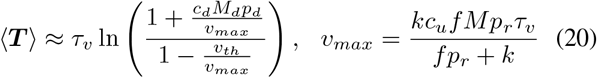

and is only defined for sufficiently large input frequencies *f* for which *v*_*max*_ *> v*_*th*_.

To quantify stochasticity in AP timing we use the following previously developed small-noise approximation

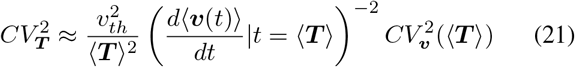

[34] –[37] where

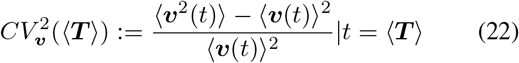

is the noise level in the membrane potential at *t* = ⟨ ***T*** ⟩. As per (21), the timing noise is proportional to noise in the voltage and inversely related to the slope squared of ⟨***v***(*t*)⟩. *Thus a flatter approach of* ⟨***v***(*t*)⟩ *to the threshold will amplify timing fluctuations*. Referring the reader to the Appendix for more details, an approximate analytical formula for 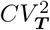 is obtained using (21) by solving the second order moment dynamics for the membrane potential ⟨***v***^2^(*t*)⟩, using equations (12) and (13).

### A. Noise in **T** in the limit k_d_ → ∞, k → ∞, p_d_ → 1

Using the approach detailed above we obtain the following result in the limit of high presynaptic refilling rate *k* → ∞ and a probability of release for the postsynaptic neuron close to one (*p*_*d*_ → 1)

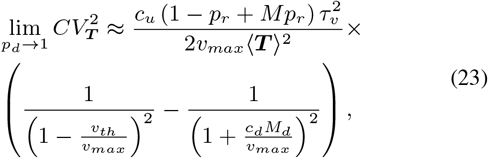

where

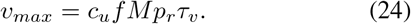

In the limit of a pure postsynaptic integrator, i.e., *τ*_*v*_→ ∞ in (10), the equations in (20) and (23) further simplify as

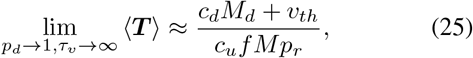

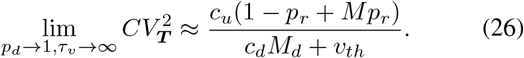

In these ideal limits, one can see from (25) and (26) that increasing autapse or negative feedback strength *c*_*d*_ elongates inter-AP times’ mean ⟨***T***⟩ and attenuates timing noise (Fig. 3).

**Fig. 3.**
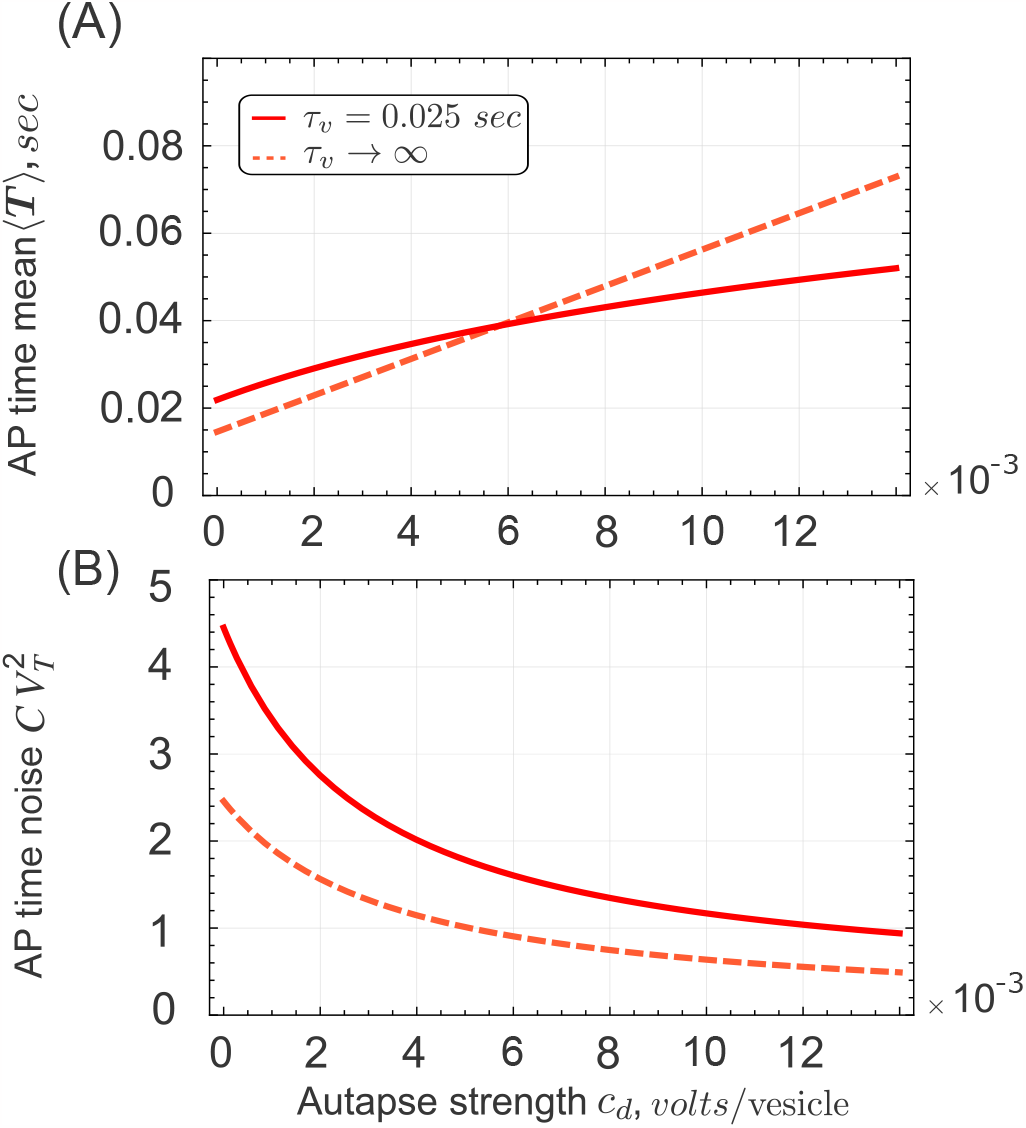
Plots of mean and noise levels, (20) and (23), respectively as a function of autapse or negative feedback strength *c*_*d*_. Increasing *c*_*d*_ lowers 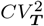 providing an effective buffering of timing stochasticity. Note that the noise levels are higher for a leaky-integrator (*τv* = 0.025 *sec*) vs. a pure-integrator *τv* → ∞. For these plots, parameters taken as *k*→ ∞, *k*_*d*_→ ∞, *c*_*u*_ = 0.02 *volts/vesicle, f* = 30 *Hz, M* = *M*_*d*_ = 20, *p*_*r*_ = 0.4, *p*_*d*_ = 1, *v*_*th*_ = 0.07 *volts* and *v*_0_ = 0.

Perhaps a better noise comparison is to keep ⟨***T***⟩ fixed with increasing *c*_*d*_ which we do in Fig. 4 by appropriately lowering the voltage-firing threshold *v*_*th*_. In this comparison, 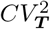 becomes invariant of *c*_*d*_ for a pure integrator (*τ*_*v*_→ ∞), but we see significant noise attenuation for a leaky-integrator (finite *τ*_*v*_). This attenuation can be intuitively linked to the negative feedback resulting in a higher slope of membrane potential buildup as compared to the open-loop case (Fig. 4A). However, for a pure-integrator where buildup is always linear, there is now no noise difference as the integration time window is kept constant by fixing ⟨***T*** ⟩.

**Fig. 4.**
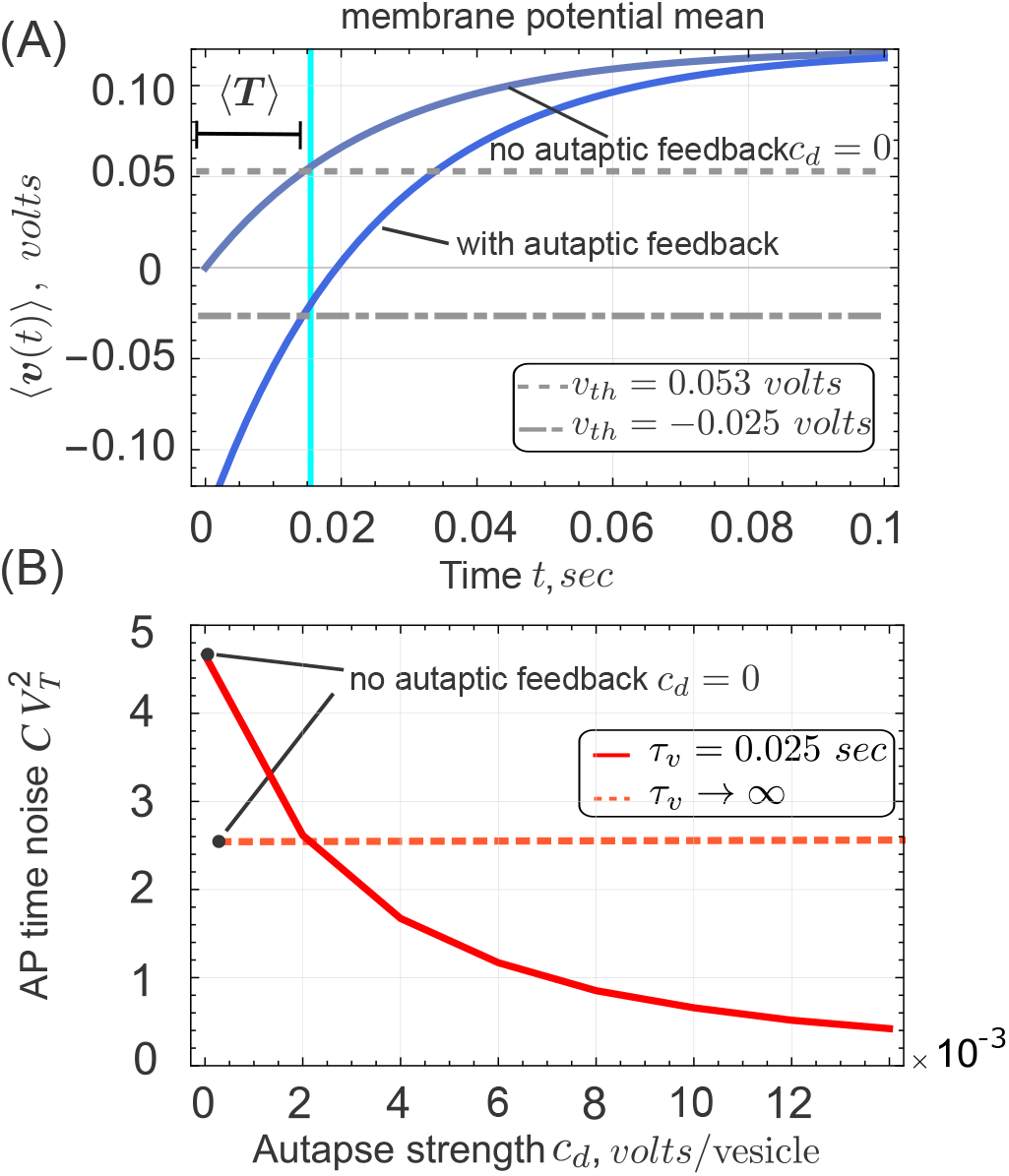
**A**. Plot of the mean potential buildup over time as given by (32) with (*c*_*d*_ = 0.007 *volts/vesicle*) and without autapse (*c*_*d*_ = 0) for *τ*_*v*_ = 0.025 *sec* and *v*_0_ = 0. The threshold (gray dashed lines) are adjusted to keep ⟨ *T* ⟩ same. **B**. Plot of the timing noise (23) as a function of *c*_*d*_ for fixed mean ⟨ ***T*** ⟩ = 0.0145 *sec*, where noise attenuation is only seen for a leaky integrator (*τv* = 0.025 *sec*). Other parameters are *cu* = 0.02 *volts/vesicle, f* = 30 *Hz, M* = *M*_*d*_ = 20, *p*_*r*_ = 0.4, *p*_*d*_ = 1.

### B. Relaxing the assumption on p_d_

The analytical results in the previous subsection were in the limit *p*_*d*_→ 1 that makes the output of the postsynaptic neuron and its voltage reset upon AP firing deterministic. For instance, ***v*** ⟼ *v*_0_ − *c*_*d*_*M*_*d*_ with probability one. Relaxing this assumption for 0 *< p*_*d*_ ≤ 1 yields

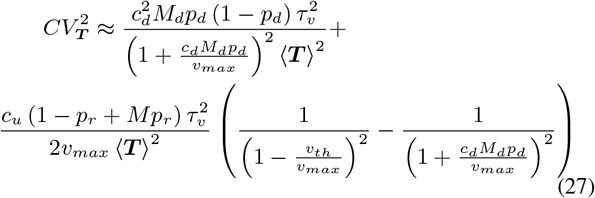

where *v*_*max*_ is in (24). A comparison with (23) shows an additional noise term on the left-hand-side of (27) that arises for *p*_*d*_ *<* 1 and captures the contribution from noise in the number of vesicles released from the autaptic axon terminal that actuates the negative feedback. In the limit *τ*_*v*_ → ∞, (27) simplifies to

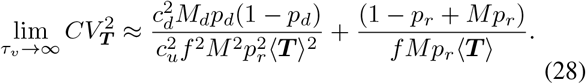

that clearly shows for fixed ⟨***T*** ⟩, 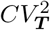 increases (remains constant) with increasing negative feedback strength *c*_*d*_ when *p*_*d*_ *<* 1 (*p*_*d*_ = 1).

### C. Stochastic simulation of the SHS model

After obtaining analytical insights we finally focus on the original SHS model (Fig. 2) relaxing prior assumptions made on *k* and *k*_*d*_ (*k*→ ∞, *k*_*d*_→ ∞). In Fig. 5 we plot the noise in the postsynaptic AP firing times with autapse normalized to its value without autapse as a function of parameters *p*_*d*_, *M*_*d*_, and *k*_*d*_.

**Fig. 5.**
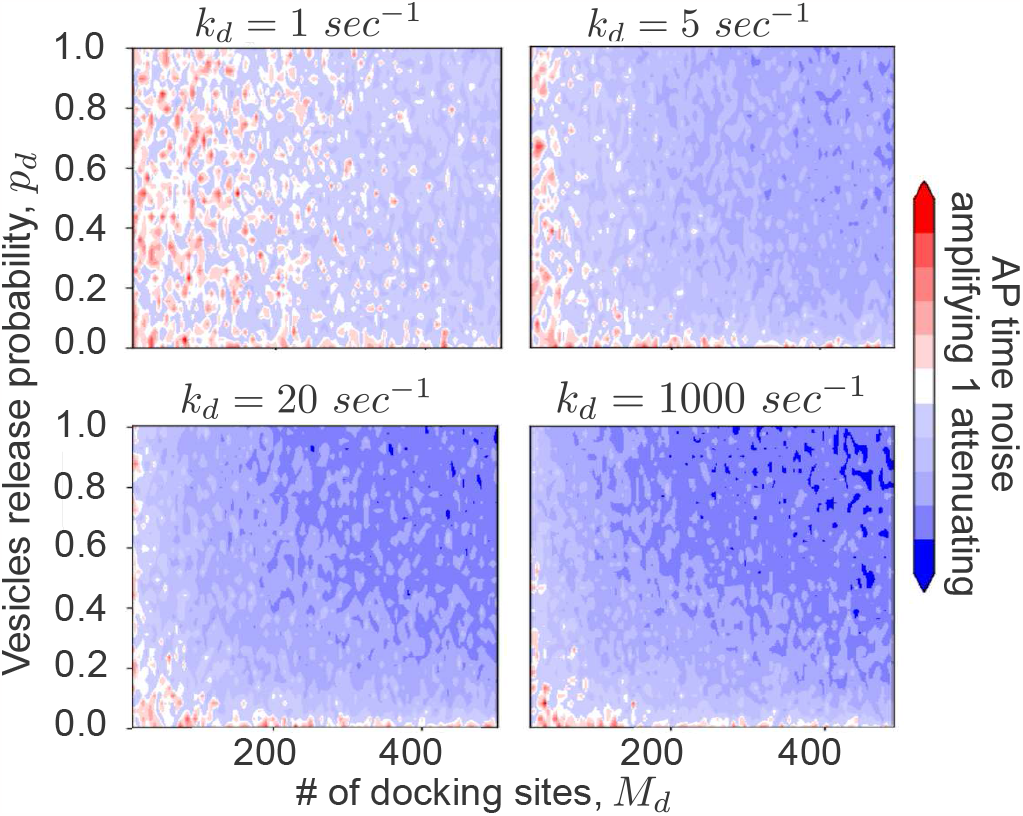
The presented heat maps depict the noise in postsynaptic AP timing *CV*_***T***_ as obtained by running a large number of stochastic simulations of the SHS model in Fig. 2 for *c*_*d*_ = 0.02. This noise is normalized by its corresponding value without autapse *c*_*d*_ = 0. The blue (red) regions present noise attenuation (amplification) where this ratio *CV*_***T***_ (*c*_*d*_ = 0.02)*/CV*_***T***_ (*c*_*d*_ = 0) is lower (higher) than one with increasing shading corresponding to a higher degree of effect. The ratio is plotted as a function of parameters *M*_*d*_, *p*_*d*_ for four different values of *k*_*d*_ with other parameters taken as *cu* = 0.02 *volts/vesicle, f* = 50 *Hz, M* = 100, *pr* = 0.3, *τ*_*v*_ = 0.01 *sec, v*_*th*_ = 0.2 *volts* and *v*_0_ = 0.

The bottom-left plot in Fig. 5 corresponds to large values of *k*_*d*_ and shows a strong noise buffering effect (blue shaded region) consistent with our analytical findings in Fig. 3 with the strongest noise buffering occurring for *p*_*d*_ = 1 and large *M*_*d*_. This buffering effect diminishes with decreasing values of *M*_*d*_ and *p*_*d*_, and a noise amplification region arises (red shaded) for sufficiently small values that is a result of noisy autaptic vesicle release. As *k*_*d*_ is lowered, the amplification region begins to dominate across all parameter values.

## IV. Conclusion

This contribution has systematically investigated the role of autaptic feedback in driving precision in postsynaptic AP firing times as has been experimentally seen [17]. We used the SHS formalism together with analytical results on FPT moments to derive analytical conditions when noise buffering occurs, and also identify regimes where timing fluctuations are amplified. At an intuitive level, negative feedback elongates the time windows for integrating bursty secretion of upstream excitatory neurotransmitter levels leading to effective noise attenuation. However, if the release of the inhibitory neurotransmitter from the postsynaptic neuron is itself quite noisy (which occurs at low values of *p*_*d*_), autapses can have the opposite effect of amplifying noise in the first-passage times.

It is interesting to point out that when keeping the postsynaptic inter-AP firing time fixed, (for example, by lowering the voltage threshold in the negative feedback case as compared to the no feedback case), we still see noise buffering; although the parameter regime corresponding to it is significantly reduced. In the latter case, when the average time to AP generation is fixed, both systems (with and without feedback) have the same integration time. Given the leaky nature of membrane potential, noise buffering here is based on lowering the threshold for the feedback case to a regime where voltage increase is linear and far away from its steady-state level. This effect has been seen in timing problems in cell biology where a protein needs to reach a critical threshold level, and timing fluctuations are minimal (for a given fixed mean FPT) in the linear regime of protein increase [35], [38], [39]. Supporting this argument, when we remove the voltage decay rate (*τ*_*v*_ → ∞) noise buffering is lost when the mean firing time is fixed (Fig. 4).

Our recent collaboration on dopamine neurotransmission has elegantly shown forms of regulation where the secreted dopamine inhibits presynaptic processes involved in the synthesis/release of the neurotransmitter [40], [41]. Mechanistically, they occur via auto-receptors on the presynaptic membrane that sense neurotransmitters in the synaptic cleft and modulate the subsequent release of vesicles [40]–[43]. As a simple example, the buildup of high levels of neurotransmitters in the synaptic cleft due to several presynaptic APs can reduce the probability of the release of synaptic vesicles to subsequent APs. We plan to investigate such types of feedback in future work and compare and contrast the noise-buffering properties of different types of feedback implementations. Incorporation of such feedback leads will lead to nonlinear SHS that we can either numerically simulate or use analytical frameworks considering small-noise regimes or moment closure schemes [44]–[47].

Several experimental works have implicated incoherent feedforward neuronal circuits [48], [49] in enhancing precision in AP firing times and these would be ripe for future analysis. Some work along these lines has been recently done for feedforward genetic circuits [50]. Finally, another interesting avenue of research is to consider autapses that form positive feedback loops as have been experimentally reported [51], and they could play a key role in optimizing signal detection.

## ACKNOWLEDGMENT

This work is supported by NIH/NIDCD grant 1R01DC019268-01 and NSF grant ECCS-1711548.

## Appendix

When all *M*_*d*_ docking sites are available for release (*k*_*d*_→ ∞) and upon the arrival of an AP at the axon terminal of the postsynaptic neuron, ***b***_*d*_ vesicles release

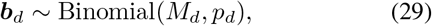

and the mean and variance of ***b***_*d*_ follows

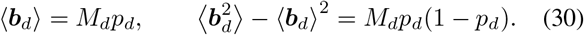

By substituting *v*_0_ = 0 and ⟨***b***_*d*_⟩ (30) in (16), we obtain

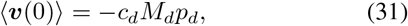

and the solution of membrane potential mean dynamics (11) follows

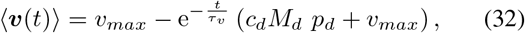

where *v*_*max*_ is as in (20). An approximate solution for the mean of ***T*** in (17) as a random variable is given by solving

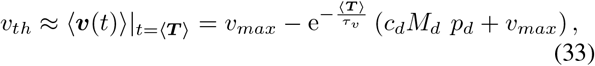

to obtain (20). Furthermore, we solve (13) for ⟨***n***(*t*)***v***(*t*)⟩ using the number of presynaptic vesicles at the steady state 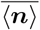 (7) and ⟨***v***(0)⟩ in (31). Therefore,

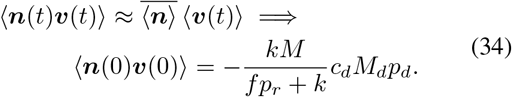

Additionally, using (16)

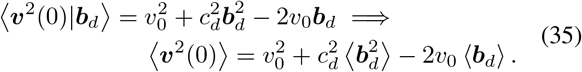

By utilizing ⟨***b***_*d*_⟩ and 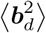 from (30) and replacing *v*_0_ = 0 into (35) yields

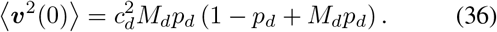

Subsequent ly, we solve (13) for ⟨***n***(*t*)***v***(*t*)⟩. Along with the steady-state mean of the docked 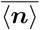 vesicles in (7) and the second order moment 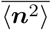 in (8), we solve (12) for ⟨***v***^2^(*t*)⟩ with ⟨***v***^2^(0) ⟩ in (36). Finally, the noise in the postsynaptic membrane potential at the mean of ***T*** follows

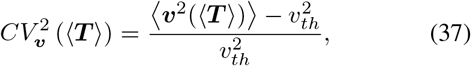

and the noise expressions in the postsynaptic AP time ***T*** follows (21) where ⟨***T*** ⟩ is in (20).

